# Targeting Tenascin-C-Toll-like Receptor 4 signalling with Adhiron-derived small molecules - a viable strategy for reducing fibrosis in Systemic Sclerosis

**DOI:** 10.1101/2025.05.23.655116

**Authors:** Thembaninkosi Gaule, Katie J Simmons, Kieran Walker, Francesco Del Galdo, Rebecca L Ross, Hema Viswambharan, Jahnavi Krishnappa, Jack Pacey, Martin McPhillie, Darren C Tomlinson, Azhar Maqbool

**Affiliations:** Clinical Population and Sciences Department, Leeds Institute for Cardiovascular and Metabolic Medicine, School of Medicine, University of Leeds, Leeds, United Kingdom; Discovery and Translational Science Department, Leeds Institute for Cardiovascular and Metabolic Medicine, School of Medicine, University of Leeds, Leeds, United Kingdom. School of Biomedical Sciences, Faculty of Biological Sciences & Astbury Centre, University of Leeds, Leeds, UK; Leeds Institute of Rheumatic and Musculoskeletal Medicine, Faculty of Medicine and Health, University of Leeds, Leeds, United Kingdom; Scleroderma Programme, NIHR Leeds Musculoskeletal Biomedical Research Centre, Leeds, United Kingdom; School of Chemistry, University of Leeds, Leeds, United Kingdom; School of Molecular and Cellular Biology, University of Leeds, Leeds, United Kingdom

**Author notes:** Joint 1st authors.

**Keywords:** Tenascin C, TLR4, Adhiron, small molecules, fibrosis, systemic sclerosis

## Abstract

Tissue fibrosis is a hallmark of systemic sclerosis (SSc) and results from the excessive production and deposition of collagen and other extracellular matrix proteins by resident fibroblasts. This excessive connective tissue accumulation leads to tissue disruption and subsequent dysfunction in the skin, lungs and other internal organs. Recent studies highlight a role for the matricellular protein Tenascin C in SSc, whereby its stimulation of Toll-like receptor 4 triggers self-sustaining fibroblast activation and ensuing fibrosis. We have utilised Adhiron guided ligand discovery to generate small molecules that target the fibrinogen-like globe domain of Tenascin C, a region involved in Toll-like receptor 4 activation and have demonstrated a reduction in the profibrotic phenotype of human dermal fibroblasts. These studies may facilitate the development of effective targeted therapy for fibrosis in SSc and other fibrotic diseases. Moreover, it highlights the utility of Adhiron guided ligand discovery to generate small molecule inhibitors to selectively modulate proteins.

**Highlights:**

- Adhiron screening was undertaken to identify binders to the FBG domain of Tenascin C and the crystal structure of the FBG-Adhiron binding was resolved.
- Adhiron guided ligand discovery was used to generate small molecule inhibitors that selectively target the FBG domain of Tenascin C.
- SPR was used to show binding of these molecules to the FBG domain of Tenascin C
- The small molecule inhibitors generated attenuated TGFβ-mediated fibrosis in dermal fibroblasts

## 1. Introduction

Persistent fibrosis a hallmark of systemic sclerosis (SSc) results from the excessive production and accumulation of collagen and other extracellular matrix (ECM) proteins by activated fibroblasts, leading to tissue disruption and dysfunction in the skin, lungs and other internal organs. Despite recent advances, clinical evidence reveals that up to one in three patients show fibrosis progression even while receiving recommended immunosuppressive or anti-fibrotic treatments [1–3] highlighting the urgent need for novel and more effective therapeutic strategies to treat Ssc.

Recent studies have implicated a role for the matricellular protein and endogenous Damage Associated Molecular Pattern (DAMP), Tenascin C (TNC), in the pathogenesis of SSc. Tenascin C is highly upregulated in both the affected tissue and circulation of SSc patients with both early and late-stage disease. It is markedly expressed in skin and lung biopsies and in blister fluid in SSc patients, and elevated serum levels of TNC correlate with the Modified Rodnan skin score, a measure of skin fibrosis [4,5]. Dermal fibroblasts explanted from SSc patients show constitutive production of TNC *in vitro*, implying that its increased accumulation in SSc may result, in part, from its cell-autonomous overproduction [4]. Moreover, treatment of healthy fibroblasts with the profibrotic cytokine TGF-β, which is strongly implicated in the pathogenesis of SSc, induces the synthesis of TNC, whereas TNC deficiency attenuates TGF-β-mediated fibrosis *in vivo* and following murine lung injury [6].

Tenascin C has been shown to elicit Toll like receptor 4 (TLR4)-dependent profibrotic responses in dermal fibroblast, including promoting myofibroblast differentiation, upregulation of αSMA and collagen expression [4]. In support of this, mice lacking TNC are protected from both skin and lung fibrosis following bleomycin injury [4, 6, 7]. Similarly, loss of either TNC or TLR4 is associated with reduced hypodermal fibrosis in the Tsk/+ mouse, a spontaneous fibrosis model [4, 8]. Importantly, the profibrotic effects of TNC are completely abrogated in TLR4-deficient fibroblasts and by small molecules inhibitors that selectively block TLR4 underscoring a key pathogenic role for TNC-TLR4 signalling in driving persistent fibrosis [9].

Collectively, these findings support a model of self-sustaining, fibroblast activation in SSc in which tissue damage causes local generation and accumulation of TNC, which then through TLR4 triggers potent stimulatory effects on fibrotic gene expression, myofibroblast differentiation, as well as enhancing the sensitisation of fibroblasts to the profibrotic effects of TGF-β [4, 10]. This persistent fibroblast activation results in enhanced matrix production, further tissue damage and TNC production, so establishing a non-resolving loop of pathological fibrosis characteristic of SSc [9]. Disrupting this non resolving loop is regarded as a viable strategy for the treatment of fibrosis in SSc.

Tenascin-C is a multi-modular protein comprising four distinct domains: an assembly domain, a series of epidermal growth factor-like repeats (EGF-L), a series of fibronectin type III-like repeats (FNIII), and a C-terminal fibrinogen-like globe (FBG) [11]. As the FBG domain of TNC has been shown to be essential for binding to and activating TLR4 [12–14], selectively preventing TLR4 activation by targeting the FBG domain represents a promising antifibrotic therapy. It is noteworthy that conventional TLR4 inhibition approaches have proven ineffective in clinical trials, and moreover, run the risk of compromising the host response to infection [15], highlighting the need for selective strategies.

The Adhiron (formally known as Affimer) technology is a novel platform based on a highly diverse library (>3×10^10^) of small, engineered binding proteins, each displaying two variable nine-amino acid loops [16]. This platform enables high affinity, selective binding to specific target proteins. Adhirons have been successfully employed to identify functional regions, probe protein function and uncover druggable pockets on previously considered undruggable proteins such as Ras [17–26]. Analogous to antibodies [27], Adhirons can also serve as valuable tools for small molecule drug discovery with the potential act as pharmacophore templates that inform design and reduce the risk of failure in early development.

In the present study we have employed Adhiron guided ligand discovery to develop small molecule that target the FBG domain of TNC. These compounds effectively attenuate the profibrotic phenotype of human dermal fibroblasts, supporting a novel strategy for small molecule development.

## 2. Material and Methods

### 2.1 Adhiron guided small molecule discovery

#### 2.1.1. Production and Purification of recombinant FBG protein

The genes encoding the FBG domains from human were synthesised by Genscript (Piscataway, USA). The FBG coding regions were amplified by PCR, digested with NheI and NotI, and cloned into a similarly digested pGEX vector modified to contain a N-terminal AviTag and C-terminal 8’His tag. The FBG domains were expressed in Shuffle cells (NEB). Single colonies were used to inoculate 5 mL of LB media with 100 μg/mL disodium carbenicillin and grown at 37 °C with shaking at 220 rpm for 16 h. The overnight culture was used to inoculate large scale cultures of 400 mL pre-warmed LB medium with 100 μg/mL carbenicillin and grown at 37 °C, and 220 rpm. When OD600 reached 0.6-0.8 expression was induced using 0.1 mM IPTG and grown for a further 16h at 25 °C with shaking at 150 rpm. Cells were harvested and resuspended in lysis buffer (50 mM NaH2PO4; 300 mM NaCl; 30 mM Imidazole; 10% Glycerol; pH 7.4) with 0.1 mg/ml of lysozyme, 1% Triton X-100, 10 U/ml Benzonase® endonuclease and 1× Halt protease inhibitor cocktail and incubated at 4°C for 1 hr. The FBG domain were purified from the supernatant using a two-step purification procedure, Ni^2+^-NTA chromatography followed by size exclusion chromatography. The supernatants were filtered through a 0.2 µm syringe filter prior to loading on a home-made Ni^2+-^NTA columns. The protein-bound columns were washed with wash buffer (50 mM NaH2PO4; 500 mM NaCl; 20 mM Imidazole; pH 7.4) to remove unbound proteins. 50mM NaH2PO4, 500 mM NaCl; 300 mM Imidazole; pH 7.4 was used to elute bound FBG. The purity of the samples was assessed by SDS-PAGE with Coomassie staining and the elution fractions were pooled accordingly. The pooled samples were loaded onto a HiLoad® 26/600 Superdex® 200 size exclusion column, eluted in PBS and concentrated using Vivaspin® centrifugal concentrators with 10 kDa cut-off.

#### 2.1.2. Phage display

Adhiron selection was performed as described previously [28, 29] using a phage display library constructed by the BioScreening Technology Group (BSTG), University of Leeds, UK. Adhiron selection was performed as described previously [28]. Briefly, biotin acceptor peptide (BAP) human Tenascin-C FBG was bound to streptavidin-coated wells (Pierce) for 1 h. Following 3h of pre-panning, the phage were incubated in wells containing human Tenascin-C FBG domain for an hour. Unbound or weak binding phage were washed away using PBST. Bound phage was eluted with 50 mM glycine–HCl (pH 2.2) for 10 min, neutralised with 1 M Tris–HCL (pH 9.1), further eluted with 100 mM triethylamine for 6 min, and neutralised with 1 M Tris–HCl (pH 7). After three biopanning rounds, 96 randomly picked positive Adhiron clones were evaluated for their binding ability to FBG by phage ELISA. Phage Elisa was undertaken in the manner we have previously described, briefly, recombinant FBG was immobilized directly on the plastic surface of 96-well Nunc MaxiSorp® plates (Thermo Scientific, Waltham, MA, US). The binding of Adhiron clones was detected with the use of 3,3’,5,5’-Tetramethylbenzidine. DNA was sent for DNA sequencing to allow analysis of the binding loop sequences.

#### 2.1.3. Production of Adhiron protein and purification

Recombinant production of the Adhiron proteins was performed as previously described [28]. Briefly, the relevant Adhiron coding regions were amplified by PCR, using Forward primer (5′-ATGGCTAGCAACTCCCTGGAAATCGAAG) and Reverse primer (5′-TACCCTAGTGGTGATGATGGTGATGC). The products were digested with NheI and NotI restriction enzymes at 37°C for 2 h and cloned into a similarly digested pET11a vector modified to contain an 8’His tag coding sequence. BL21 STARä (DE3) cells were transformed using the relevant cloned and cultured as described in Tiede et al. [28]. The cells were harvested and resuspended in phosphate buffered saline pH 7.4 supplemented with 0.1 mg/ml of lysozyme, 1% Triton X-100, 10 U/ml Benzonase® endonuclease and 1× Halt protease inhibitor cocktail and incubated at 4°C for 1 hr. Adhirons were purified from supernatant using Ni^2+^-NTA affinity chromatography as previously described [28].

#### 2.1.4. Solid Phase inhibition assays

Streptavidin coated 96-well plates were blocked overnight at 4 °C using 2 × blocking buffer (Sigma) in PBS-T (0.1% Tween-20). 5 μg/ml of Bap tagged hFBG-C was immobilised onto the pre blocked streptavidin coated 96 well plates for 1 hr at room temperature. Unbound Bap tagged hFBG-C was washed off using PBS-T. 25 μg/ml of Adhiron was then added to appropriate wells and incubated for 1 hr at room temperature with gentle agitation. Excess Adhiron was removed by washing three times with PBST. To assess the Adhiron protein’s ability to block the hFBG-C/TLR4 25 μg/ml of TLR4 was added to the wells and incubated for 2 hr at room temperature. TLR4 binding was detected as described previously [30]. After three washes TLR4 binding was visualised by adding 50 μl of TMB and measuring absorbance at 620 nm. Experimental controls involved probing for TLR4 binding in the absence of Adhiron.

#### 2.1.5. Crystallization, data collection, and structure determination

8’His tagged human FBG binding Adhiron and untagged human FBG were expressed in BL21 DE3 star and shuffle cells respectively. The cells were harvested by centrifugation at 10 000 g for 20 mins. The pellets resuspended in PBS supplemented with benzonase, triton X-100, and lysozyme were incubated at room temperature with gentle agitation for 1 hr. The 8’His tagged Adhiron protein was immobilised onto Ni-NTA beads by incubating the Adhiron protein containing lysate with Ni-NTA agarose beads for 1 hr at room temperature. Following the 1 hr incubation the Ni-NTA beads were washed to remove unbound Adhiron protein. The Ni-NTA beads carrying Adhiron were then added to the human FBG containing lysate and incubated for 1 hr at room temperature. The beads were washed and the Adhiron/human FBG complex was eluted using PBS supplemented with 300 mM imidazole. The contents of the elutions were analysed using SDS-PAGE and the appropriate elutions were pooled, concentrated, and further purified using and size exclusion column (HiLoad® 26/600 Superdex® 200). The Adhiron/human FBG complex was concentrated to 30 mg/ml and crystallised using the sitting drop method and the MCSG-3 crystal screen (molecular dimensions). The crystal belonged to space group P 21 21 21 with cell dimensions a=77.570 Å,b=90.140 Å and C=104.410 Å and a=b=g=90. The Adhiron/human FBG structure was determined using molecular replacement.

#### 2.1.6. Ligand-Based Virtual Screening

A shape similarity screen was performed using OpenEye’s ROCS software [31]. ROCS allows rapid identification of active compounds by shape comparison, through aligning and scoring a small molecule database against a query structure of residues N41, H43 and D47 from Adhiron 52. The EON software compares electrostatic potential maps of pre-aligned molecules to a query structure, producing a Tanimoto similarity score for comparison. This was used in conjunction with the eMolecules library of 10,000,000 commercially available small molecules. Identification of compounds best matching the shape and electrostatic properties of the Adhiron residues interacting with FBG was carried out using the graphical user interface Maestro [32] and the OpenEye graphical user interface VIDA looking for specific hydrogen bonding interactions with FBG and minimising any steric clashes with the protein. This work was undertaken on ARC4, part of the High-Performance Computing facilities at the University of Leeds. Molecules were selected for biological evaluation based on favourable shape and electrostatic matching to the query residues.

#### 2.1.7. Metabolic Stability and Solubility Assessment

This was carried out by Eurofins Ltd according to their standard protocols.

#### 2.1.8. Surface plasmon resonance

A Biacore 1K (Cytiva, USA) was used to analyse the interaction between fibrinogen and a small molecule. Briefly, biotinylated fibrinogen was first diluted to 1 μM with 200 μL of 1x phosphate buffered saline (pH 7.4) + 0.1% tween and then flowed across the surface of a SA sensor chip (Cytiva, USA) at a flow rate of 30 μL/min, reaching ∼ 6400 response units (RU). All the binding experiments were performed at 25°C at a continuous flow rate of 30 μL/min with 1x phosphate buffered saline (pH 7.4) + 0.1% tween + 1% DMSO to maintain compound solubility. Due to the use of DMSO and its high refractive index contribution, solvent correction was applied, and signal corrected against the control surface response. Serial concentrations of the small molecules were run across the chip surface at 7.8125, 15.625, 31.25, 62.5, 125 and 250 μM. All the equilibrium constants (KDs) used for evaluating binding affinity were determined with Biacore Insight Evaluation Software (Cytiva, USA).

### 2.2 Experimental protocols

All participants provided written informed consent according to a protocol approved by Medicine and Health Regulatory agency (NRES-011NE to FDG, IRAS 15/NE/0211). The privacy rights of the human subjects have been observed. All SSc patients fulfilled the ACR/EULAR 2013 criteria [33]. Patients were included in an at-risk population if they presented with Raynaud’s and any very early diagnosis of SSC (VEDOSS) criteria [34–36]. By definition, at-risk patients did not meet ACR/EULAR 2013 classification criteria for SSc (score <9), neither did they fulfil classification for any other connective tissue disease.

#### 2.2.1. Cell culture

Full thickness skin biopsies were surgically obtained from the forearms of three adult healthy controls (HC), three adult patients with recent onset SSc, defined as a disease duration of less than 18 months from the appearance of clinically detectable skin induration. All patients satisfied the 2013 ACR/EULAR criteria for the classification of SSc and had diffuse cutaneous clinical subset (dcSSc) as defined by LeRoy et al [37], and 6 very early diagnosis of SSc (VEDOSS) patients. Fibroblasts were isolated from skin biopsies by expansion out of the cut biopsies, and primary cell lines established after two passages. Human telomerase-immortalisation was carried out using retroviral transduction as described [38,39]. Experimental testing on dermal fibroblast primary cell lines were performed within 10 passages. Cells were maintained in Dulbecco’s modified Eagle medium (DMEM) (Gibco) supplemented with 10% FBS (Sigma) and penicillin-streptomycin (Sigma). TGF-β1 treatment (10 ng/ml) was used in 1% FBS conditions following an overnight starvation in 1% FBS.

#### 2.2.2. Western blotting

Total proteins were extracted from fibroblasts in RIPA buffer and resolved by SDS-PAGE (10-15% Tris-Glycine). Proteins were transferred onto Hybond nitrocellulose membranes (Amersham biosciences) and probed with antibodies specific for TNC (Santa Cruz Biotechnology) and α-smooth muscle actin (Abcam) or loading control β-actin. Immunoblots were visualized with species-specific HRP conjugated secondary antibodies (Sigma) and ECL ThermoFisher Scientific (/Pierce) on a Biorad chemiDoc imaging system.

#### 2.2.3. Quantitative Real time PCR

RNA was extracted from cells using the RNA extraction kit (Zymo Research) following the manufacturing protocols. RNA was reverse transcribed using the cDNA synthesis kit (ThermoFisher Scientific). Q-RT-PCR were performed using SyBr Green PCR kit (ThermoFisher Scientific) with primers specific for *TNC* (Forward 5’-TCGACGTGTTTCCCAGACAG-3’; Reverse 5’-AACGGTGTCTTCCAGAGCAG-3’), *COL1A1* (Forward; CCTCCAGGGCTCCAACGAG Reverse; TCTATCACTGTCTTGCCCCA), *COL1A2* (Forward; GATGTTGAACTTGTTGCTGAGC Reverse; TCTTTCCCCATTCATTTGTCTT), *ACTA2* (Forward; TGTATGTGGCTATCCAGGCG Reverse; AGAGTCCAGCACGATGCCAG), *CCN2* ((Forward; GTGTGCACTGCCAAAGATGGT Reverse; TTGGAAGGACTCACCGCT) and *GAPDH* (Forward; ACCCACTCCTCCACCTTTGA Reverse; CTGTTGCTGTAGCCAAATTCGT). The data obtained was analysed according to the ΔΔ Ct method. *GAPDH* served as housekeeping gene. Composite profibrotic score was calculated using the mean value of ΔΔ Ct of *COL1A1, COL1A2, ACTA2* and *CCN2* for each sample.

#### 2.2.4. Collagen gel contraction assay

Collagen gel contraction assays were prepared using Cell Contraction Assay Kits (Cell Biolabs), per manufacturer instructions. Briefly, 2x10^5^ fibroblasts were cultured within collagen gel for 16 hr at 37°C 5% CO2, then released from the sides of wells and photos taken over 48 hr. The percentage change in gel area relative to area of gel at 0h was analysed with ImageJ software.

#### 2.2.5. Live Dead Cell Assay

Fibroblasts were seeded at a density of 20,000 cells per well in a 24 well plate. Cells were treated with DMSO vehicle or small molecule (10μM) for 24 hr. Experiments were performed in triplicate. Cell viability was determined using LIVE/DEAD™ fluorometric assay according to the manufacturer’s instructions (Invitrogen).

#### 2.2.6. Statistical analysis

Categorical variables were presented as numbers and percentages, while continuous variables were reported as mean ± standard deviation (SD), mean ± standard error (SEM), or median with interquartile range (IQR) depending on the data distribution. Comparisons between groups were conducted using the student t-test or chi-square test. Statistical significance was defined as a p-value less than 0.05 for all analyses, and all tests were two-tailed. Data analysis was performed using RStudio (version 2023.03.0) or GraphPad Prism software (version 9.5.1).

## 3. Results

### 3.1 TNC mRNA and protein levels elevated in SSc and VEDOSS dermal fibroblasts correlates to levels of profibrotic gene expression

TNC gene expression was observed to be higher in fibroblasts isolated from SSc patients as well as in fibroblast from VEDOSS patients, being approximately 7-fold and 5.5-fold greater than HC levels for SSc and VEDOSS respectively (**Figure 1A**), however due to heterogeneity in the different donor SSc fibroblast cell lines, this did not reach significance. When assessing the profibrotic gene expression of the biological triplicates of HC and SSc dermal fibroblast cell lines, by assessing mRNA levels of *CCN2, COL1A1, COL1A2*, and *ACTA2*, all SSc fibroblasts samples collectively showed increased gene expression compared to HC, with significant differences observed with *COL1A1* and *COL1A2* (**Figure 1B**). Combining these genes into a profibrotic score shows SSc fibroblasts have an increased level compared to HC (**Figure 1C**). Interestingly, there is a strong correlation with the profibrotic score and TNC gene expression in all fibroblasts cell lines, showing a direct link between the fibrotic phenotype of SSc dermal fibroblasts and TNC expression (**Figure 1D**). Moreover, all single genes show strong correlation with TNC expression (Spearman Correlation R>0.7986, P<0.0027), with *CCN2* showing the strongest correlation (R=0.9142 P<0.0001). Moreover, the SSC3 cell line with low basal TNC expression, had the lowest *CCN2* expression (**Figure 1B and C**) which is in line with previous studies that demonstrate an induction of TNC following CCN2 stimulation [40–42]. We also demonstrate that TNC expression is enhanced by profibrotic cytokine, TGF-β1, in HC dermal fibroblasts (**Figure 1E**). TGF-β1 stimulation of 24 hr triggered an increase in fibroblast TNC expression by 6-fold (**Figure 1E**). This is consistent with the observations of Bhattacharyya who demonstrated a dose- and time-dependent TGF-β-induced upregulation of TNC in both neonatal and adult skin fibroblasts [4].

**Figure 1:**
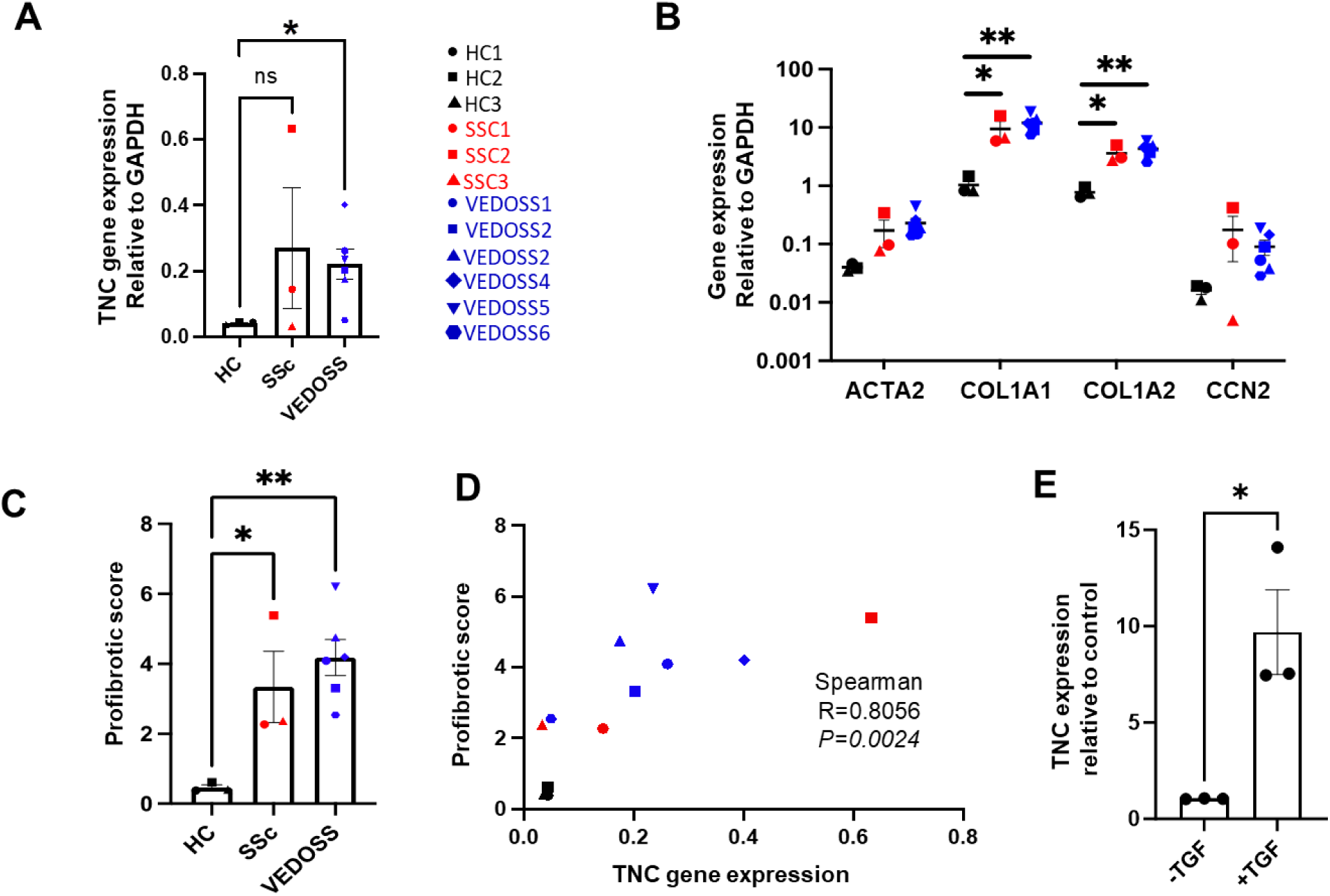
TNC expression in dermal fibroblasts correlates to levels of profibrotic markers. TNC mRNA expression in HC (n=3), SSc (n=3) and VEDOSS (n=6) fibroblasts in starved conditions **(A)**. Each symbol represents a cell line generated from a single donor (HC; Black, SSc; Red, VEDOSS; Blue). Gene expression analysis for *ACTA2, COL1A1, COL1A2, CCN2* relative to housekeeping gene *GAPDH* for all dermal fibroblast cells line in A **(B)**. Combination of analysis from B into a profibrotic score for each category of dermal fibroblast **(C)**. Correlation between TNC gene expression and profibrotic score for data in B and C **(D)**. TNC mRNA expression in HC dermal fibroblasts (n=3) with and without TGF-β1 **(E).** Bar charts represent mean ±standard error (SE). Statistical tests used; unpaired student t-test (two tailed) (A, B, C and E) and spearman correlation (D). Statistical significance illustrated; *=P<0.05, **=P<0.01 and ns = non-significant.

### 3.2 Adhiron guided small molecule identification

#### 3.2.1 Identification and Isolation of Adhiron with inhibitory properties

Adhirons have been previously shown to bind to ‘hot spots’ on protein surfaces blocking protein function through steric and allosteric mechanisms [18, 21, 22, 43]. Here we determined the ability to isolate Adhirons capable of blocking the interaction of the FBG domain of TNC (FBG-C) with TLR4. The BAP tag labelled FBG-C domain was expressed in shuffle cells and purified using a two-step purification method (**Figure 2A**). In vivo biotinylation of the FBG-C domain allowed isolation of Adhiron reagents by phage display. Biotinylation was confirmed by western blotting and ELISA (**Figure 2B**). The purified bap tagged FBG-C domain was panned against the Adhiron phage library [28, 44, 16]. After three panning round 96 clones were picked and assessed using phage ELISA [28].

**Figure 2:**
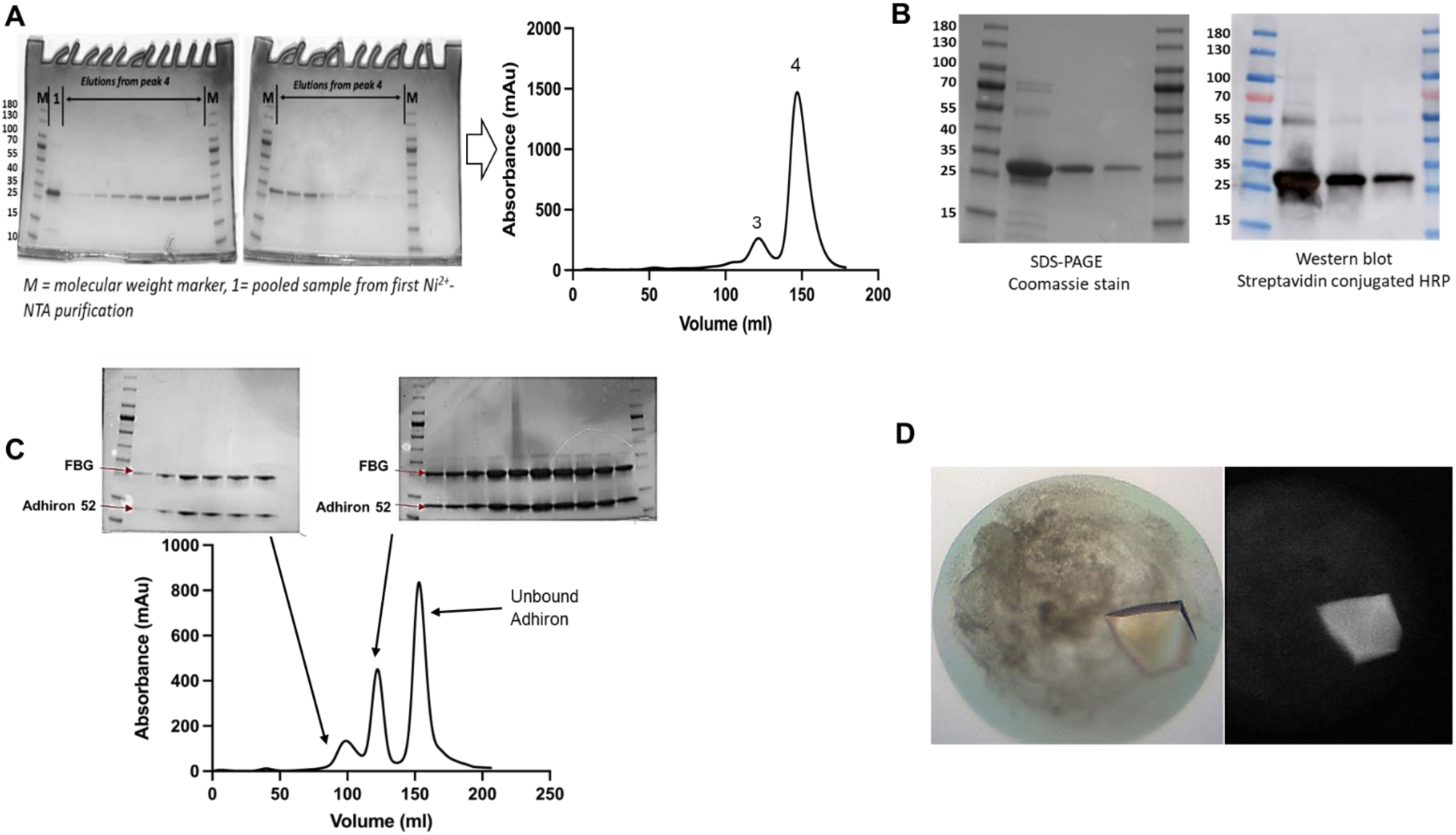
Purification and crystallisation of the FBG-Adhiron 52 complex. Two step purification of the recombinant FBG protein using Ni2+-NTA chromatography followed by size exclusion chromatography. Fractions from peak 4 were run on the gel (**A**). Protein purity assessment by SDS-PAGE with Coomassie staining -SDS-PAGE gel showing purified his-tagged and avi -tagged FBG from human TNC (right panel), western blot of Avi tagged FBG probed using streptavidin HRP (left panel) (**B**). Purification and crystallisation of the FBG-Adhiron 52 complex, denaturing SDS PAGE gels of fractions from the size exclusion columns (**C**) and crystal of the FBG-Adhiron 52 complex (**D**).

Seven out of the 96 Adhirons were non-specific as they interacted with both FBG-C domain and the control (**Figure 3A**). DNA sequencing highlighted a consensus motif on variable region I across the 86 clones. Due to factors such as expression yields, solubility, and stability, six Adhirons were taken forward for further characterisation. The six Adhirons were subcloned into pET11a, expressed in BL21 DE3 star cells and purified. The Adhirons were tested for the ability to block the TLR4/FBG-C interaction. The solid phase inhibition assay showed that as expected the scaffold (control Adhiron) did not inhibit the TLR4/FBG-C interaction. At the same concentrations, Adhirons 9, 12, 32, 45 and 52 showed different degrees of inhibition whereas Adhiron 47 did not inhibit (**Figure 3D**). Variable region 1 of Adhiron 47 did not have the conserved motif like Adhirons 9, 12, 32, 45 and 52 suggesting that Adhiron 47 may have a different epitope compared to other Adhirons (**Table 1**). The solid phase inhibition assays showed that Adhiron 52 displayed the best inhibition (**Figure 3D**) and SPR indicated binding affinities of 15.3nM and 11.8nM for human and mouse FBG-C, respectively (**Figure 3B, C**).

**Table 1:**
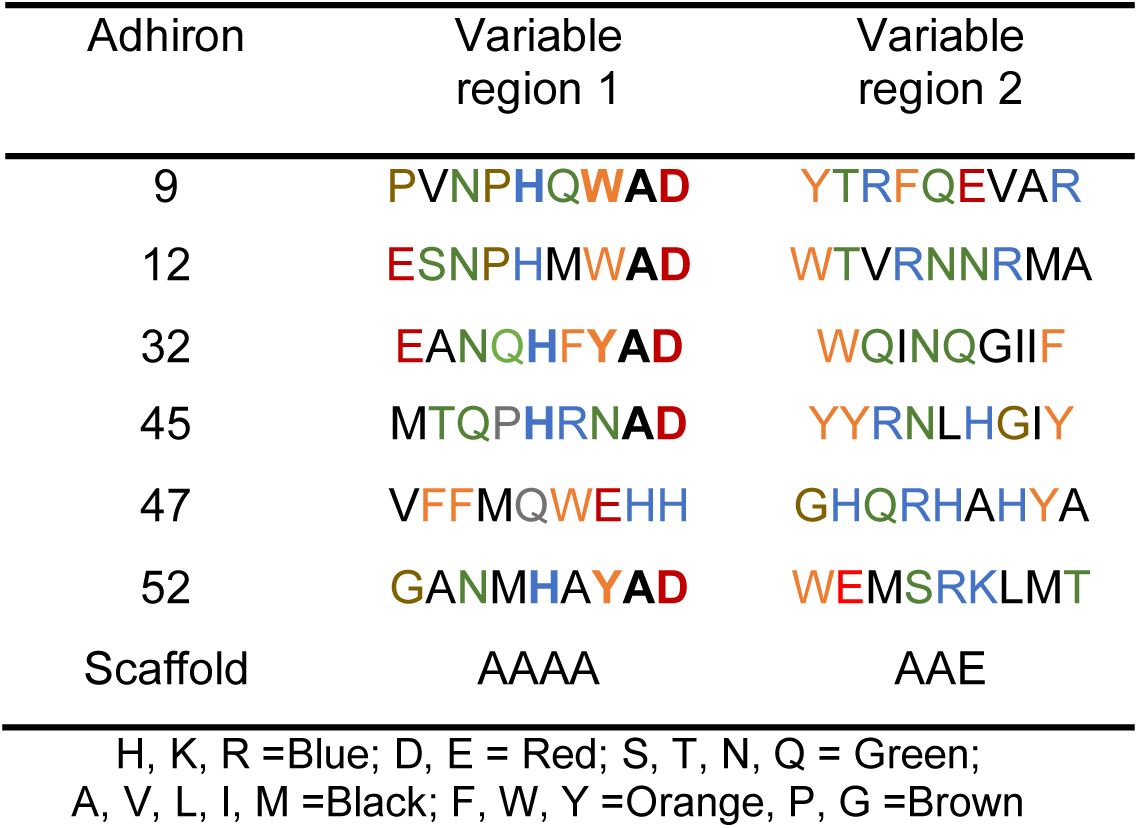
Adhiron loop sequences.

**Figure 3.**
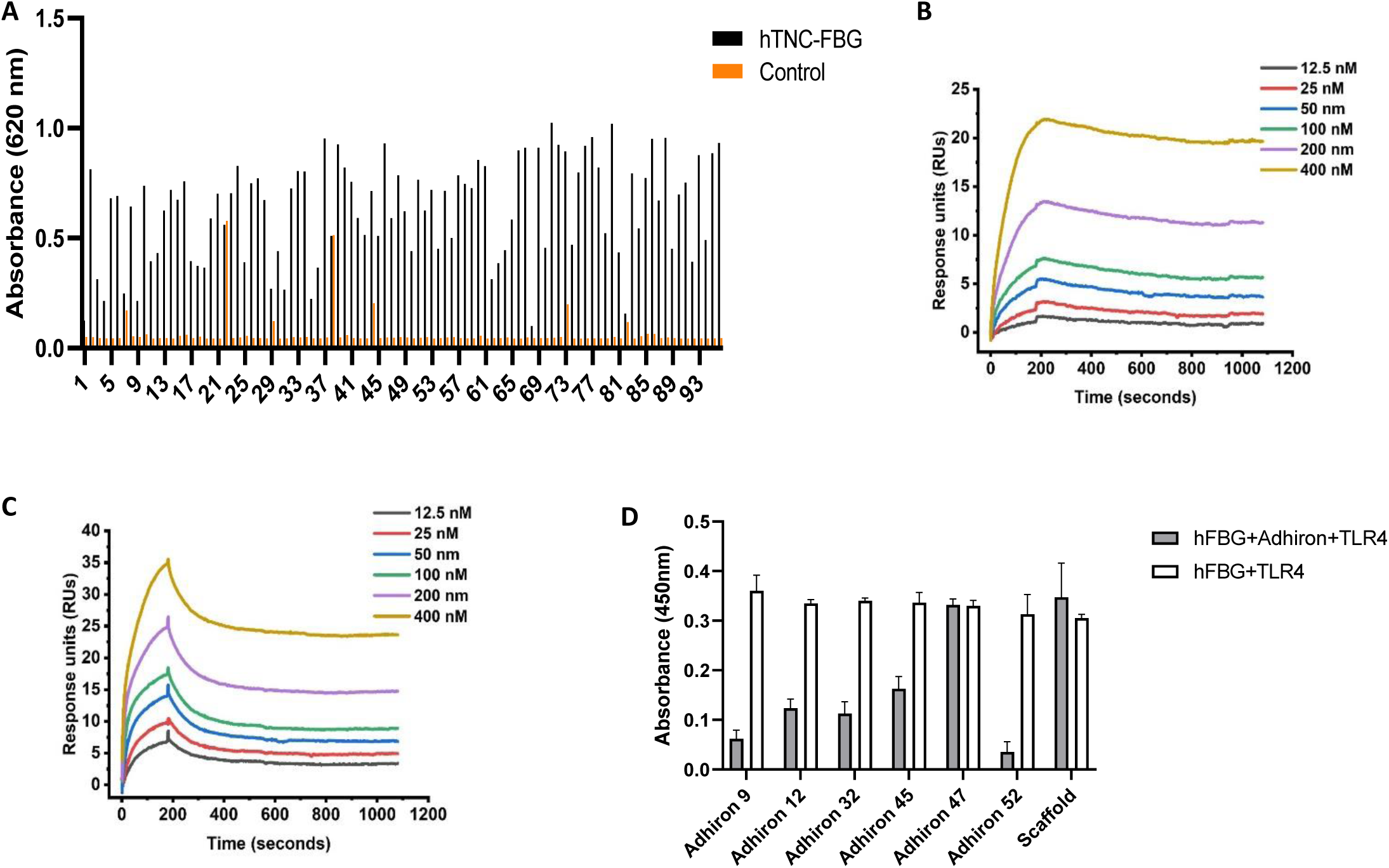
Phage ELISA and solid phase inhibition assay: Phage ELISA results from 96 Adhirons identified in screens against the FBG domain of human TNC **(A**). Kinetic analysis of Adhiron 52 binding to human **(B)** and mouse **(C)** FBG-C by surface plasmon resonance (SPR). Representative sensorgram obtained from injection of Adhiron at concentrations of 12.5, 25, 50, 100,200 and 400nM. Solid phase inhibition assay. 96-well plates coated with 5 µg ml−1 of FBG-C (Human) were incubated with TLR4 only (white) and TLR4 in the presence of Adhiron (grey). Data are shown as mean ± SEM, n = 3 (**D**).

#### 3.2.2. Structural insights into Adhiron 52 and FBG-C domain interaction

To gain atomic insights into how Adhiron 52 interacts with FBG-C domain, the Adhiron/FBG-C complex was produced and crystallised (PDB ID 9R5Y, **Figure 2C and D**) in space group P21 21 21 with two complexes in the asymmetric unit. The structure was determined by molecular replacement and refined at resolution 1.4 Angstroms (**Supplementary Table 1**). In complex with Adhiron 52, the overall structure of the FBG-C domain does not change. It maintains the three subdomain ABP structure observed in FRePs (**Figure 4A**). Adhiron 52 interacts with the FBG-C at the p subdomain (**Figure 4A**). However, in comparison to uncomplexed FBG-C domain (PDB 6QNV), the Adhiron results in subtle changes in the cation ridge region, the proposed TLR4 binding site [30]. On interaction with the Adhiron, part of the cationic ridge β - strand becomes less structured and the side chains of residues E2104 and V2103 take up different conformations (**Figure 4B**). Moreover, the disordered binding loop due to the absence of Ca^2+^ observed in FBG-C structure (PDB - 6QNV) is structured in Adhiron/FBG-C complex indicating that Adhiron 52 stabilises the disordered Ca^2+^ binding loop in the absence of calcium (**Figure 4B**).

**Figure 4:**
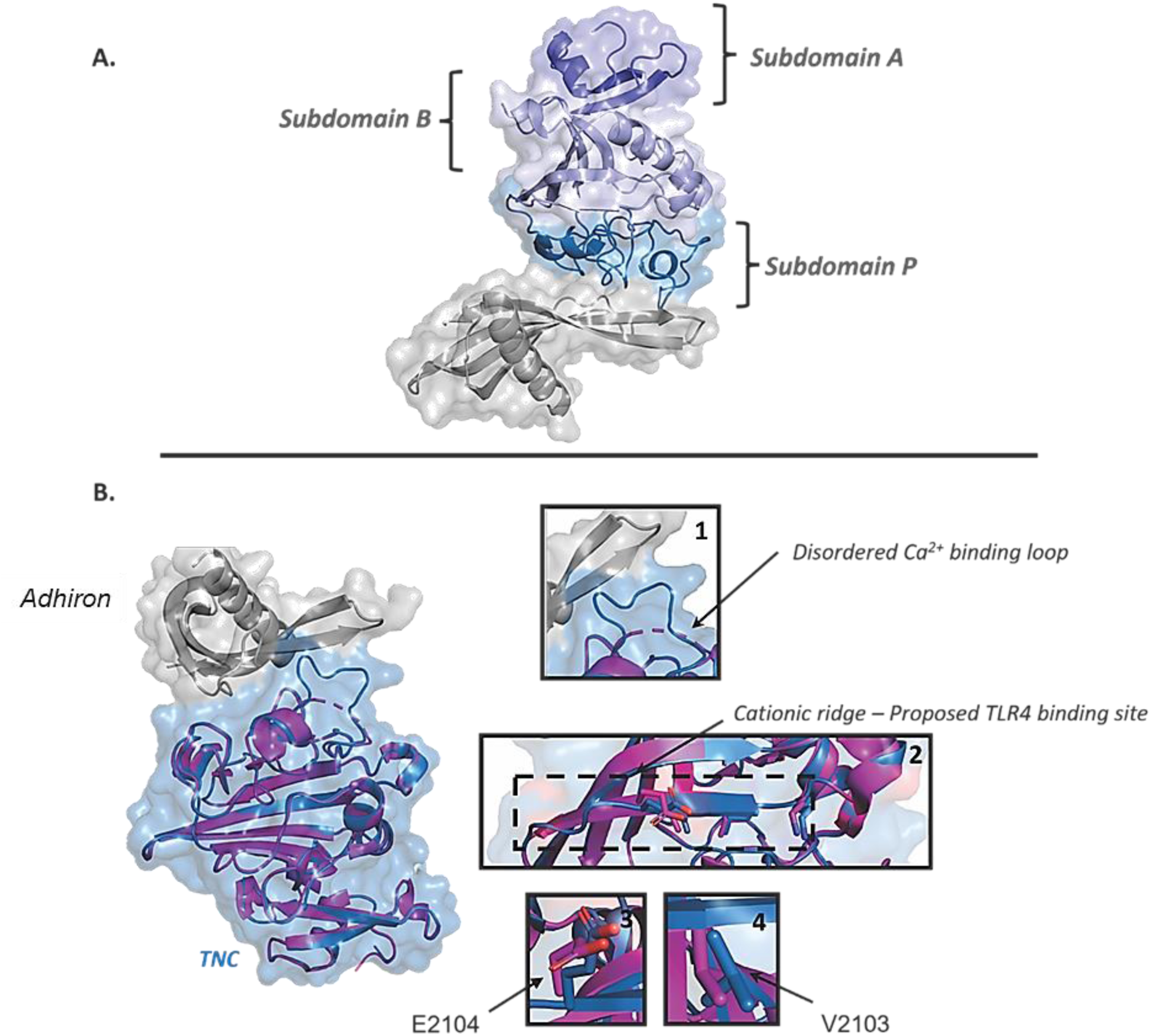
The crystal structure of Adhiron 52/FBG-C complex. A) The different shades of blue show the ABP subdomains. Subdomain A is shown in slate, subdomain B in light blue and subdomain P in sky-blue **(A**). Structural alignment of FBG-C/Adhiron 52 complex (Skyblue; PDB: 9R5Y) with non-complexed FBG-C (Magenta; PDB: 6QNV). *Insert 1* shows the structural changes in the disordered Ca^2+^ binding loop*. Insert 2* displays the subtle changes to the TLR4 binding site (cationic ridge). *Inserts 3 and 4* show conformational changes of residues E2104 and V2103 (**B**).

The crystal structure showed that variable region 1 on Adhiron 52 forms the major contact site with residues G39, N41, H43, Y45 and D47 involved in hydrogen bonds with residues H2175, H2150, S2164, Y2140, N2148 and Y2116 in subdomain P of FBG-C (**Figure 5**, inserts 1, 2 and 3). The main chain carbonyl group of G39 interacts with a hydrogen donated by a nitrogen on the imidazole group of H2175. N41 forms a hydrogen bond network where the side chain amide group interacts with the side chain hydroxyl group of S2164 while side chain carbonyl group interacts with backbone amide group of H2150. The imidazole group nitrogen atoms on H43 interact with hydroxyl group of Y2140 and the side chain carbonyl group of N2148 through hydrogen acceptor (N…H-O) and hydrogen donation (N-HN…H-O) respectively. Simultaneously, the side chain amide group of N2148 interacts with the main chain carbonyl group of Adhiron residue Y45. Adhiron residue D47 forms the final hydrogen bond network with residues Y2116 and Adhiron residue W73. The side chain hydroxyl group of D47 interacts with Y2116 while the carbonyl group interacts with the main chain amide group of Adhiron residue W73 also forms a hydrogen. Fewer hydrogen bonds were observed in the variable region 2 of the Adhiron 52 and FBG-C interaction interface (**Figure 5**). A water molecule acts as a bridge between Adhiron residue E74 and FBG-C residue R2147 (**Figure 5**, insert 6). Adhiron residues, Y43, W73, M75 and M80 stack against each other to form a hydrophobic pocket which I2133 on FBG-C snugly fits in (**Figure 5**, insert 5) and stabilises the disordered Ca^2+^ binding loop. The main chain amide group of residue M75 interacts with the carbonyl backbone group of FBG-C residue S2131 further increasing the stability of the disordered loop in the absence of calcium.

Finally, a lack of cross reactivity of Adhiron 52 with other members of the Tenascin family, notably, Tenascin R and Tenascin X was demonstrated (**Supplementary** Figure 1 and 2).

**Figure 5.**
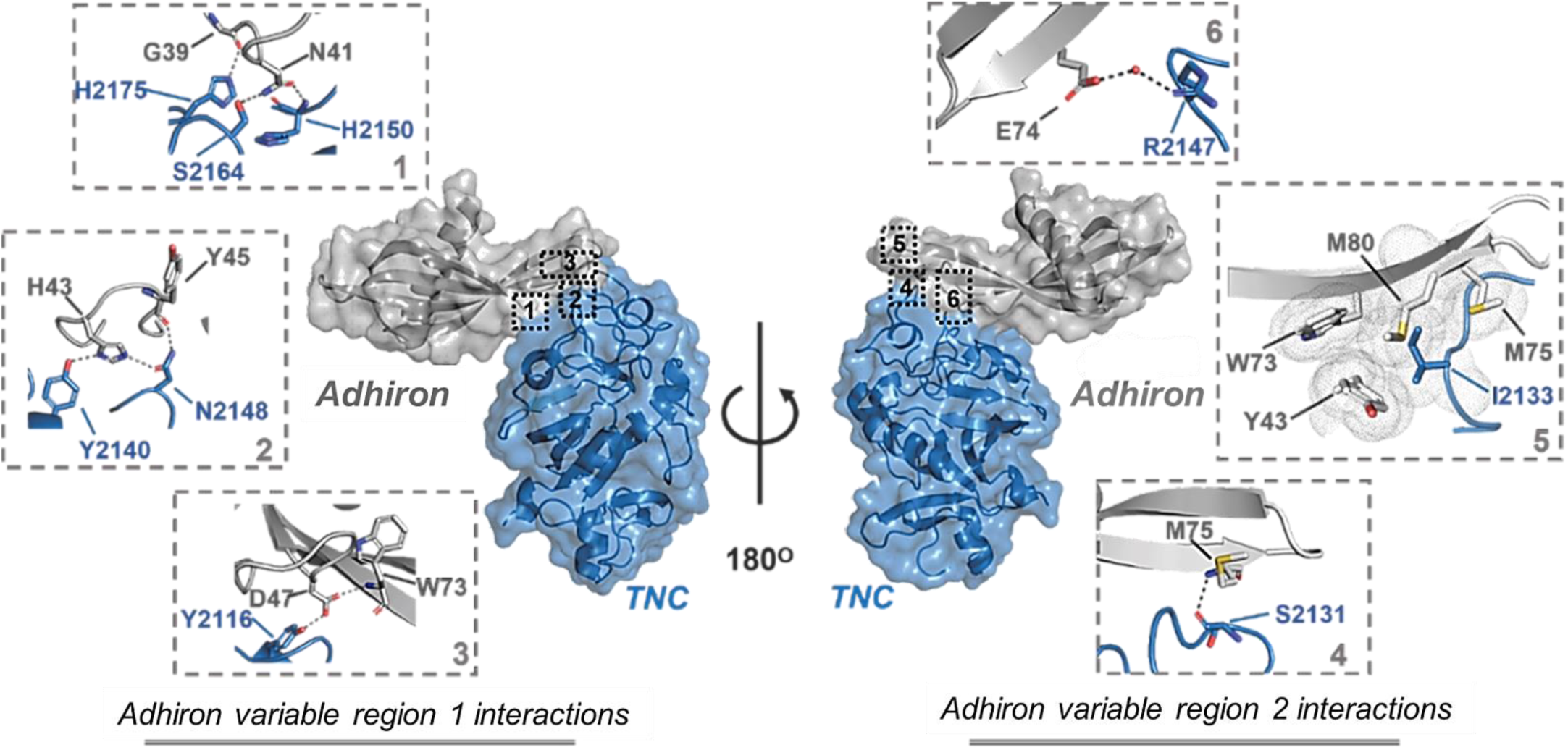
The crystal structure of FBG domain of human TNC (Skyblue) bound to Adhiron 52 (grey) (PDB: 9R5Y) inserts 1 - 3. show how five residues on the variable loop 1 of Adhiron 52 (G39, N41, H43, Y45 and D47) interact with six residues on hFBG (Y2116, Y2140, N2148, H2150, S2164 and H2175). Inserts 4 – 6 show the interactions between variable loop2 and FBG-C domain. Adhiron residues E74, Y43, W73, M80 and M75 interact with residues R2174, I2133 and S2131 on FBG-C.

#### 3.2.3. Identification of small molecules to mimic Adhiron 52

Three amino acid residues were found to be crucial for the Adhiron-FBG-C interaction: N41, H43 and D47. We used the ligand-based screening tool ROCS and identified compounds from the eMolecules library which mimicked the shape and electrostatic interactions of the key amino acid residues from Adhiron 52 (**Figure 6A-C**). Ten hit compounds were purchased (**Supplementary Table 2**) and their integrity was checked by LCMS before *in vitro* evaluation.

**Figure 6:**
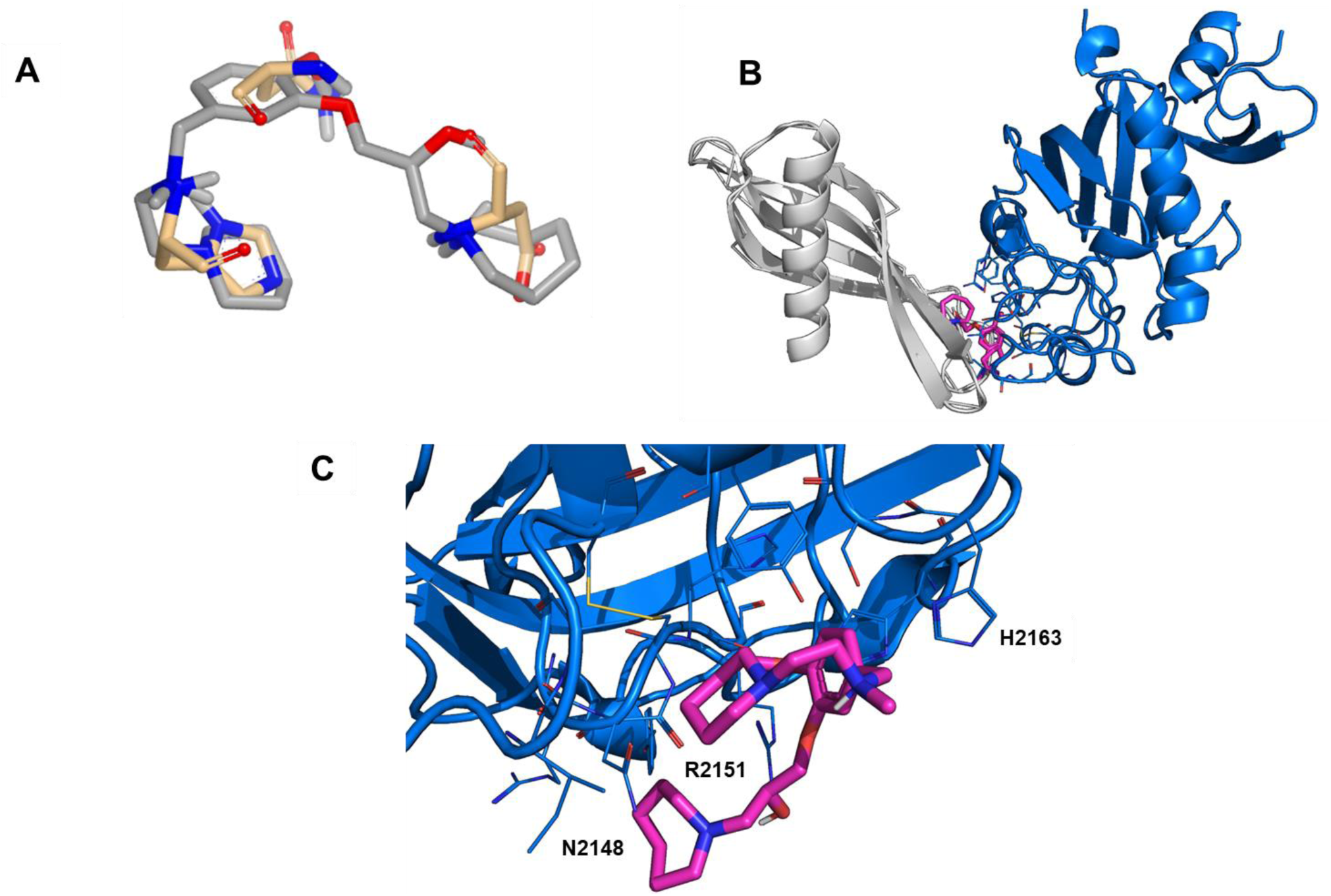
Identification of compounds as mimics of Adhiron 52. Small molecule 73494830 (grey sticks) mimics the shape and electrostatic interaction of Asn41 and His43 from Adhiron loop1 (beige sticks) **(A)**. Adhiron (grey ribbons) bound to FBG domain of TNC (blue ribbons) with predicted binding mode of 73494830 shown as pink sticks **(B).** Close up of predicted binding mode of 73494830 with the FBG domain of TNC **(C).**

### 3.3. Targeting FBG-C domain with small molecules reduces the profibrotic phenotype of human dermal fibroblasts

To test the ability of the small molecules, to modulate the interaction of the FBG-C domain of Tenascin C with TLR4, we measured their ability to reduce profibrotic responses in healthy human (HC) dermal fibroblasts. Cell lines were treated for 24 hr with 10 μM of each small molecule along with the profibrotic cytokine, TGF-β1, and the combined gene expression (expressed as a profibrotic score) of the profibrotic genes αSMA, COL1A1, COL1A2 and CCN2 was compared relative to that induced by TGF-β1 and DMSO vehicle (control, CTR) only (**Figure 7A**). Three molecules; 404, 464 and 830 reduced the TGF-induced profibrotic score relative to TGF-β1 (**Figure 7A, B**). These had no effect on cell viability as assessed using a Live/Dead cell assay (ThemoFisher Scientific, (**Figure 7C**). Due to their structural similarity these molecules were classified into 2 series: [404-series 1] and [464 and 830-series 2]. Using Surface Plasmon Resonance (SPR) we determined that compound 404 (series 1) appears to bind to the FBG domain of TNC in a non-specific manner (**Figure 7D**). However, the dissociation constant of 830 from series 2 was measured at 553 µM (**Figure 7D**).

**Figure 7:**
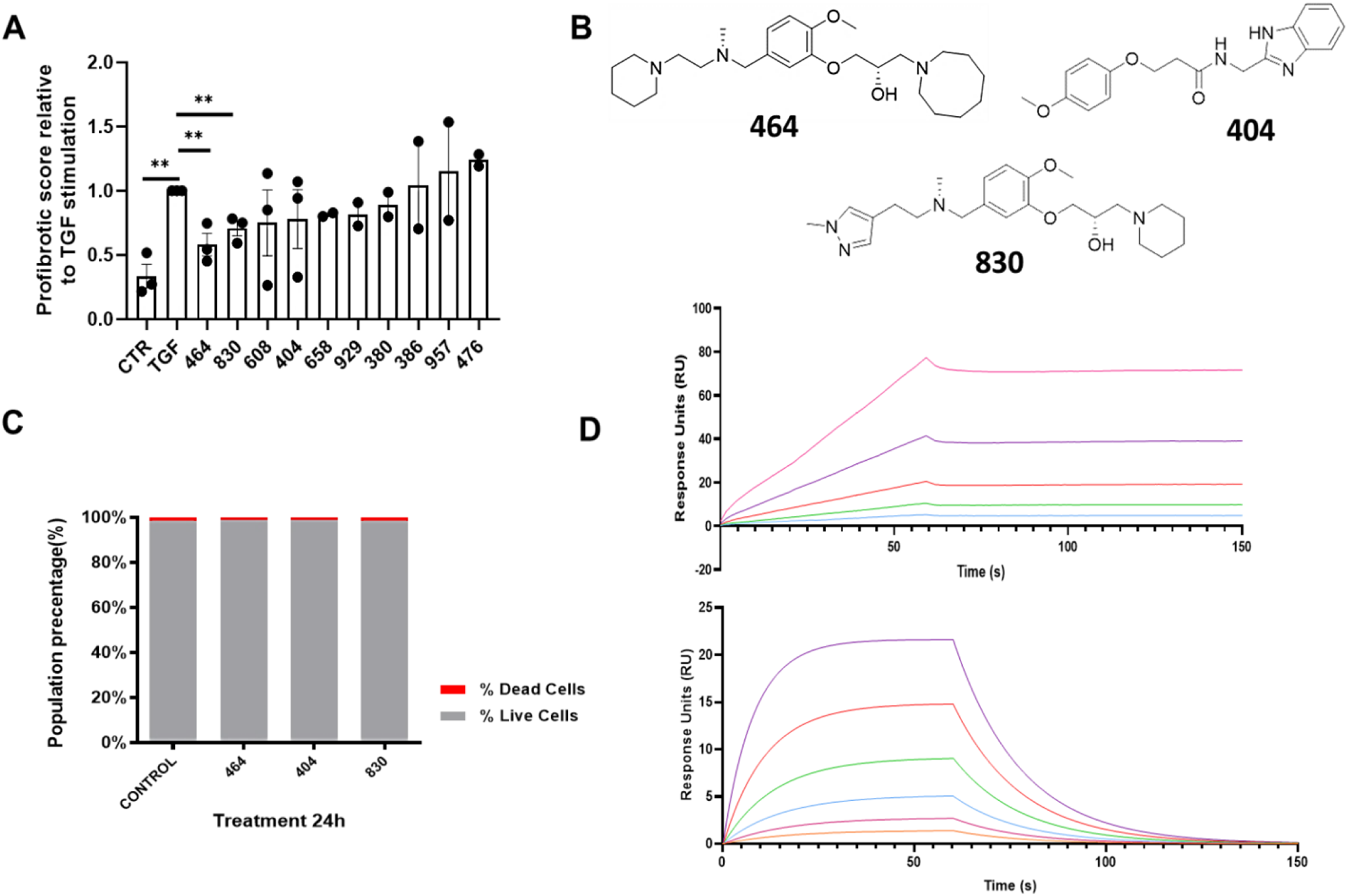
The properties of the lead candidate molecules (464, 404 and 830) on fibrosis, cell viability and FBG-C binding. The effect of inhibitors on TGF-β1 induced profibrotic score of human dermal fibroblasts. Healthy dermal fibroblasts (n=2-3, each symbol represents a cell line generated from a single donor) treated with DMSO vehicle or inhibitors in TGF-β1 conditions for 24 hrs, as well as control (CTR) conditions. Gene expression of *ACTA2, COL1A1, COL1A2* and *CCN2* relative to *GAPDH* analysed and combined into a profibrotic composite score relative to that induced in TGF-β1+DMSO only (TGF) condition **(A).** Chemical structure of compounds 464, 404 and 830 **(B)**. Effect of small molecule 464, 404, 830 (10μM, 24 hr) on fibroblast viability using a Live/Dead cell assay **(C).** Kinetic analysis of 404 and 830 binding to FBG-BAP using surface plasmon resonance (SPR) technology. **Upper panel:** Representative sensorgram of dose response obtained from injection of 404 at concentrations of 7.8125, 15.625, 31.25, 62.5, 125 and 250μM. **Lower panel:** Representative sensorgram of dose response obtained from injection of 830 at concentrations of 7.8125, 15.625, 31.25, 62.5, 125 and 250μM. Mean Kd of 533μM (n=3) **(D).**

Early stage physicochemical profiling of 404, 464 and 830 to ascertain compound metabolic stability and aqueous solubility *in vitro* demonstrated that both series displayed good solubility (>75µM) with series 2 having better human liver microsomal stability (clearance of 5-10µL/min/mg for series 2 compared to 70 for series 1) (**Table 2**).

**Table 2:** Early stage physicochemical profiling of 404, 464 and 830.

| Compound ID | Aqueous Solubility μM (pH 7.4) | Microsomal Stability (human) $CL_{int}$ (μL/min/ mg protein) |
| --- | --- | --- |
| 464 - (49306464) | >100 | <8.6 |
| 404 - (Z26782404) | >100 | 70.0 |
| 830 - (73494830) | 75 | 5 |

Due to the specific binding affinities for series 2 to the FBG domain of TNC, inhibitors 464 and 830 were retested in on dermal fibroblast isolated from SSc patients, where they were shown to reduce the composite pro-fibrotic gene score back to control unstimulated levels (**Figure 8A**). Moreover, an attenuation of the TGF-β1 induced fibrotic response by the selected inhibitors was further demonstrated in collagen gel contraction assays using HC dermal fibroblasts. With 2 hr of TGF-β1 stimulation, a significant gel contraction was induced, since gel area contracted to 85% of 0 h condition compared to CTR (97%) (**Figure 8B-C**). Treatment with inhibitor 464 (94%) and 830 (91%) were able to reduce TGF-β1 triggered gel contraction to that not significantly different to CTR levels (97%) compared to 0 hr timepoint (**Figure 8B-C**).

**Figure 8:**
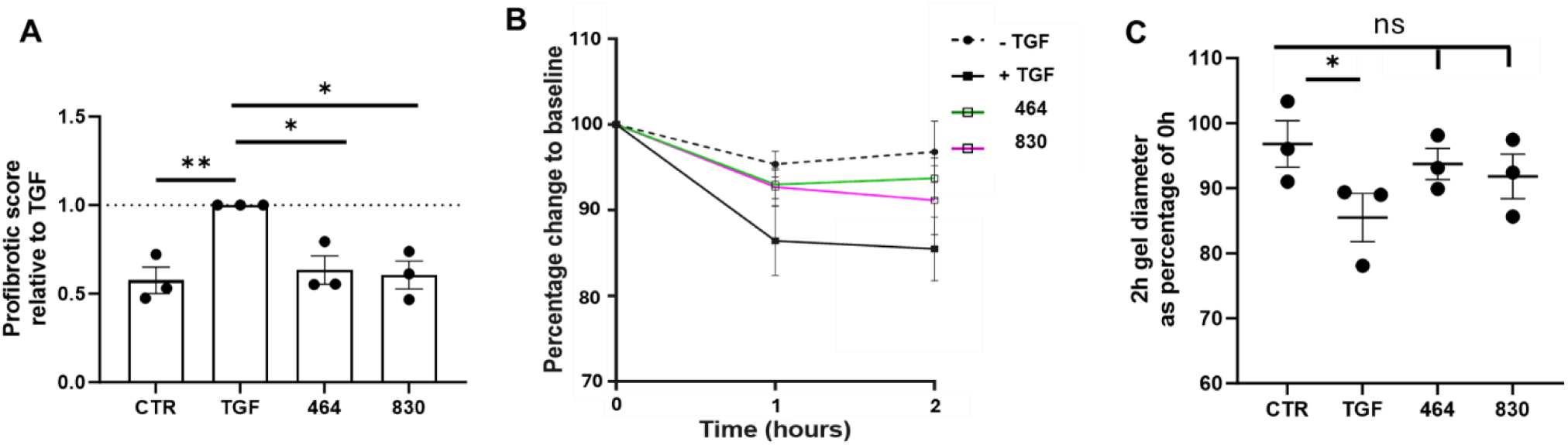
FBG-C targeting by 464 and 830 reduces the TGF-β1 induced profibrotic phenotype of human dermal fibroblasts. Reduced subset of inhibitors was tested on SSc dermal fibroblasts (n=3) and fibrotic score analysed as in Figure 7 (above) **(A).** Percentage change in gel contraction of gel cultured with healthy fibroblasts over 2 hr compared to baseline area (0 hr). Cells were treated with TGF-β1 and DMSO (+TGF), or inhibitor (464 or 830) or in control conditions (-TGF) (**B-C**).

## 4. Discussion and conclusion

### 4.1. Tenascin-C levels are increased in tissue of patients with SSc and correlate with markers of fibrosis

Constitutive production of TNC was observed in dermal fibroblasts isolated from patients with SSc compared to healthy controls. This is consistent with the current data and previous clinical observations associating high circulating TNC levels with markers of SSc and implies that the increased TNC accumulation may result from its cell autonomous overproduction by activated fibroblasts [4] (Figure 1A). Increased expression of TNC was also correlated with an increase in fibrotic gene score (notably, *ACTA2, COL1A1, COLl1A2* and *CCN2* **Figure 1B-D**). TNC has been reported to elicit cell specific responses including type 1 collagen synthesis where it drives the fibrotic response in dermal and lung fibroblasts and contributes to the integrity of the ECM in young skin [4, 45]. Moreover, TNC silencing results in the reduction of CCN2 in the kidney mesangial cells [46] and reciprocally CCN2 promotes deposition of several ECM proteins, including collagen, fibronectin and TNC [47]. Interestingly, we report for the first time that TNC expression is also raised in dermal fibroblasts isolated from VEDOSS individuals (**Figure 1A**), which have been characterised previously to show biomarkers seen in SSc fibroblasts, although from sites of absent clinical fibrosis [39]. Studies have reported a potential role for TNC in the early diagnosis of rheumatoid arthritis (RA), where the detection of antibodies to citrullinated TNC in patients with early synovitis was associated with the development of RA [48]. The early detection of TNC in dermal fibroblasts biopsies of VEDOSS patients might offer similar opportunities in identifying individuals at risk of developing SSc.

### 4.2. Tenascin C expression is induced by the profibrotic cytokine TGF-β

We demonstrated that TNC expression in HC fibroblasts is induced by the profibrotic cytokine TGF-β, this is consistent with earlier reports by Bhattacharyya *et al*. who also suggested that autonomous TNC upregulation by fibroblasts in various fibrotic disorders maybe due to autocrine stimulation by TGF-β [4]. TGF-β has been identified as a regulator of pathological fibrogenesis in SSc and related fibrotic disorders. Dysregulation of TGF-β signalling is evident in SSc and high levels of this profibrotic cytokine and its regulated genes have been detected in skin biopsies, being positively correlated with the severity of the disease [4]. Although the inhibition of TGF-β, has been demonstrated as an efficient strategy for reducing fibrosis in preclinical models, translating these findings into clinic has proved difficult due to adverse effects stemming from TGF-β’s physiological actions in inflammation and tissue homeostasis. Fresolimumab, a TGF-β neutralizing antibody, does exert anti-fibrotic effects in SSc patients, though this is accompanied by a high incidence of keratoacanthomas, which limits its use in treatment [49]. Consequentially, alternative treatments are being sought.

TGF-β increases TNC expression in cardiac [50], lung [51], intestinal subepithelial [52] and dermal fibroblast [4]. The latter was supported by our current observations, which demonstrated a 3 and 5 fold increase in TNC expression for SSc and HC fibroblasts, respectively. Previous evidence that TGF-β Receptor Kinase inhibitor SB-431542, abolished the induction of TNC by TGF-β and that TGF-β failed to induce TNC expression in Smad3-/-fibroblasts support a role for TGF-β signalling in TNC induction [4].

It is noteworthy that TNC deficiency attenuates TGF-β-mediated fibrosis in both isolated dermal and lung fibroblasts. TGF-β stimulation of type 1 collagen and αSMA was attenuated in fibroblast isolated from TNC-/-null mice compared to wild type fibroblasts [4,6]. TNC deficiency also attenuated TGF-β-mediated fibrosis following murine lung injury [6]. TNC-null mice had 85% less lung collagen than wild-type mice, following bleomycin injury (which is dependent on canonical TGF-β signalling), with the lung interstitium containing fewer myofibroblasts and cells with intranuclear Smad-2/3 staining, suggesting impaired TGF-β activation or signalling [6] and implying a role for TNC in TGF-β’s fibrotic response.

Recently a role for TLR4 signalling in the augmentation of TGF-β responses in SSc was proposed [45]. Elevated expression of TLR4, together with increased accumulation of endogenous TLR4 ligands (notably TNC, fibronectin-EDA and hyaluronic acid), in lesional skin and lung tissues from patients with SSc and in mice with bleomycin-induced SSc was reported [53]. Moreover, TLR4 activation also induced stimulation of ECM gene expression in explanted skin fibroblasts and significantly enhanced their ability to mount a profibrotic response when challenged with TGF-β1. A model for fibrogenesis was thus proposed whereby tissue damage resulting from chronic injury stimulates the local generation and accumulation of TNC, this then activates TLR4 signalling in resident fibroblasts resulting in enhanced ECM production and TGF-β secretion and a self-amplifying vicious cycle of fibrosis [9] (**Figure 9**). Targeting the TNC-TLR4 interaction represents a promising anti-fibrotic therapy. However, targeting TLR4 directly to break this feedback loop may compromise the host immune defence. Disrupting persistent TLR4 signalling by targeting the endogenous ligand’s, notably TNC, interaction with TLR4, does however, represents a potential novel strategy for breaking the cycle of progressive fibrosis in SSc [4, 9].

**Figure 9:**
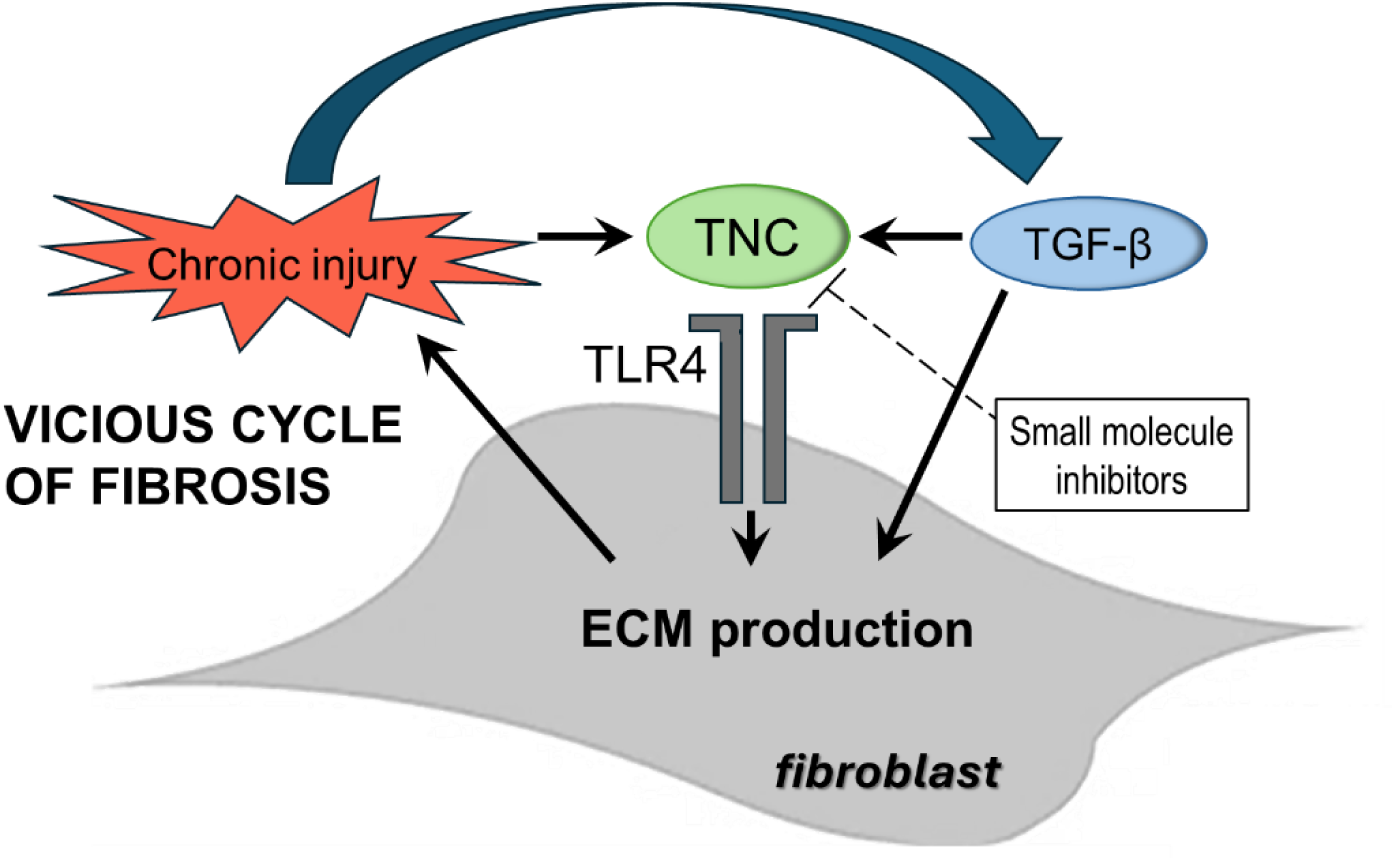
Mechanism of self-sustaining fibroblast activity and fibrosis in SSc, whereby tissue injury and TGF-β induce TNC expression resulting in the activation of TLR4 causing further tissue damage and ECM production. Adapted from [9, 54]

### 4.3. Small molecule modulators of the interaction between to the FBG domain of TNC and TLR4 can attenuate TGF-β induced fibrotic responses

We demonstrate that small molecule inhibitors 464 and 830 that target the FBG domain of TNC to block its interaction with TLR4, were able to attenuate the profibrotic effects of TGF-β. These inhibitors were designed using an Adhiron guided ligand discovery technique. Small molecules were selected based on their ability to mimic the structure of the interacting residues from the Adhiron that binds to FBG and prevents its interaction with TLR4. Specific binding of the selected inhibitors to the FBG domain was confirmed using SPR. Adhiron reagents are small non-antibody binding proteins that have been used in broad applications including in the modulation of protein function and protein– protein interactions. Although, their use in multiple protein capture-based assays (immunoassays and diagnostics) has been established, *in vivo* therapeutic application is at present in its early stages, hence the interest in developing a series of small molecule to mimics their effect.

Both 464 and 830 have structural similarity (**Figure 7B**) in that they both contain the same central substituted dimethoxyphenyl core, however 464 contains a saturated piperidine ring on the western portion of the molecule whereas 830 contains an aromatic N-methylpyrazole. The eastern portion of the molecules both contain saturated ring systems, with 830 having a piperidine whereas 464 contains the larger azapane ring. Importantly both inhibitor series display favourable ADME properties making them suitable for further optimisation (**Table 2**). Both inhibitor series were effective in reducing the expression of TGF-β stimulated fibrotic genes including αSMA, COL1A1, COL1A2 and CCN2 as well as myofibroblast differentiation as assessed by attenuation of collagen gel contraction. In conclusion, these finding indicate that small molecule selectively targeting the interaction of the FBG domain of TNC with TLR4 abrogated fibrotic responses in SSc fibroblasts in vitro following TGF-β stimulation, with 464 and 830 being promising candidates for further development as anti-fibrotic drugs. Moreover, it highlights the utility of Adhiron guided ligand discovery to generate small molecule inhibitors to selectively target protein-protein interactions.

## CRediT authorship contribution statement

**Thembaninkosi Gaul**e: Methodology, Investigation, Data curation, Writing-review and editing. **Katie J Simmons:** Methodology, Investigation, Data curation, Supervision, Writing – review & editing. **Kieran Walker:** Investigation, Data curation. **Francesco Del Galdo**: Writing-review and editing. **Rebecca L Ross**: Methodology, Investigation, Data curation, Supervision, Writing – review & editing. **Hema Viswambharan**: Investigation, Data curation. **Jahnavi Krishnappa**: Investigation. **Jack Pacey**: Investigation. **Martin McPhillie**; Investigation, Supervision. **Darren C Tomlinson**: Methodology, Investigation, Data curation, Supervision, Funding acquisition. **Azhar Maqbool**: Conceptualization, Methodology, Investigation, Data curation, Project administration, Writing – review & editing, Writing – original draft, Funding acquisition.

## Acknowledgements

This work was supported by the British Heart Foundation Project Grant No: PG/17/72/33255. Biosample collection was funded by SRUK grant and Kennedy Trust Program Foundation Grant and supported by National Institute for Health and Care Research Biomedical Research Centre (NIHR BRC) (NIHR213331) and the Susan Cheney Scleroderma Research Foundation, the Medical Research Council (MR/S001530/1) (RR, FDG, KW).

## Abbreviations

ACTA2: actin alpha 2 (alpha smooth muscle actin)
cDNA: complementary DNA
CIF: cumulative incidence function
CLIA: Clinical Laboratory Improvement Amendments
CMRI: cardiac magnetic resonance imaging
COL1A1: Collagen Type I Alpha 1 Chain
COL1A2: Collagen Type I Alpha 2 Chain
CCN2: Cellular Communication Network Factor 2 (Connective Tissue Growth Factor)
DAMP: damage associated molecular pattern
dc-SSC: diffuse cutaneous systemic sclerosis
ECL: Electrochemiluminescence
ECM: Extracellular Matrix
EGF-L: epidermal growth factor-like
EULAR: ACR European League Against Rheumatism/American College of Rheumatology
FBG: fibrinogen-like globe
FBG-C: fibrinogen-like globe of Tenascin C
FBS: fetal bovine serum
FNIII: fibronectin type III-like
FVC: forced vital capacity
GAPDH: glyceraldehyde 3-phosphate dehydrogenase
HRCT: High resolution computer tomography
HRP: horse radish peroxidase
ILD: interstitial lung disease
lcSSc: limited systemic sclerosis
MAP: Multi-Anayte Profiling
mRSS: modified Rodnan skin score
PCR: polymerase chain reaction
PFT: pulmonary function test
Q-RT-PCR: quantitative reverse transcription polymerase chain reaction
RHC: right heart catherisation
ROCS: Rapid Overlay of Chemical Structures
RNA: ribonucleic acid
SDS-PAGE: sodium dodecyl-sulfate polyacrylamide gel electrophoresis
SSc: systemic sclerosis
TGFβ: Transforming Growth Factor β
TLR4: Toll-like receptor 4
TNC: Tenascin C
VEDOSS: very early diagnosis of systemic sclerosis.

## Appendix A. Supplementary data

Supplementary data to this article can be found online at https://

## Data Statement

Data will be available on request

## Supplementary Data

**Supplementary Figure 1:**
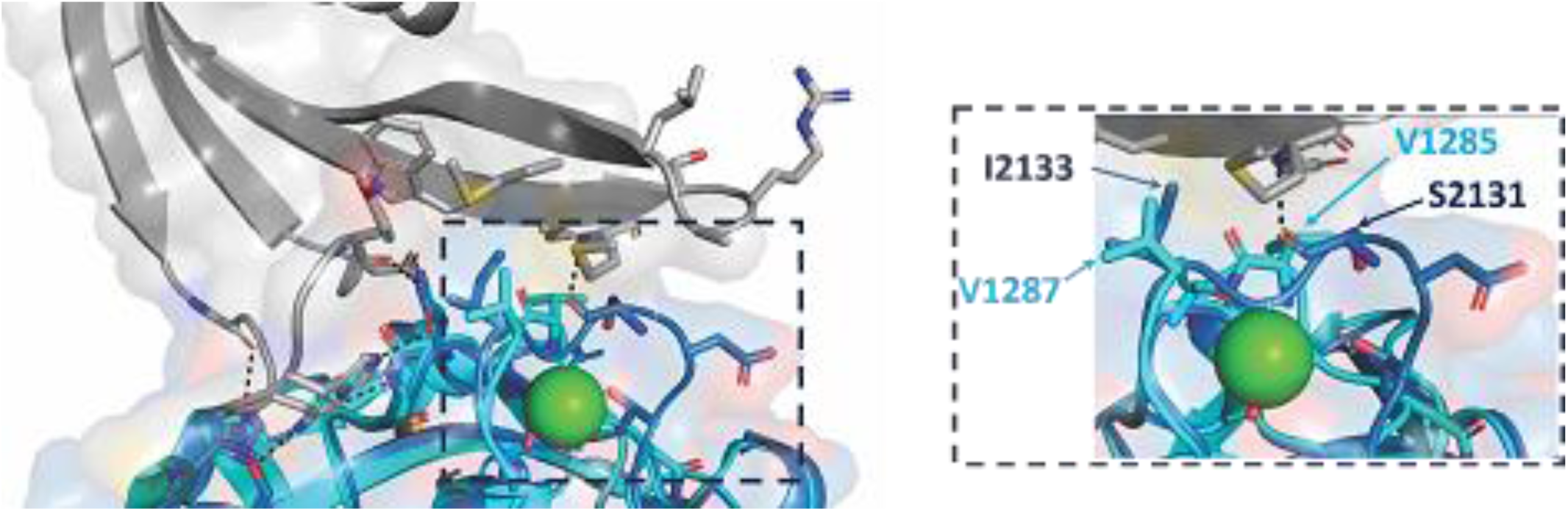
TNR-FBG (Cyan; PDB:8FNA) and Adhiron 52 interaction (Skyblue; PDB:9R5Y). **An** 2+ overlay of the TNC/Adhiron structure with the FBG domain of Tenascin R reveals differences in in Ca binding sites that would impact Adhiron 52 binding to TNR.

**Supplementary Figure 2:**
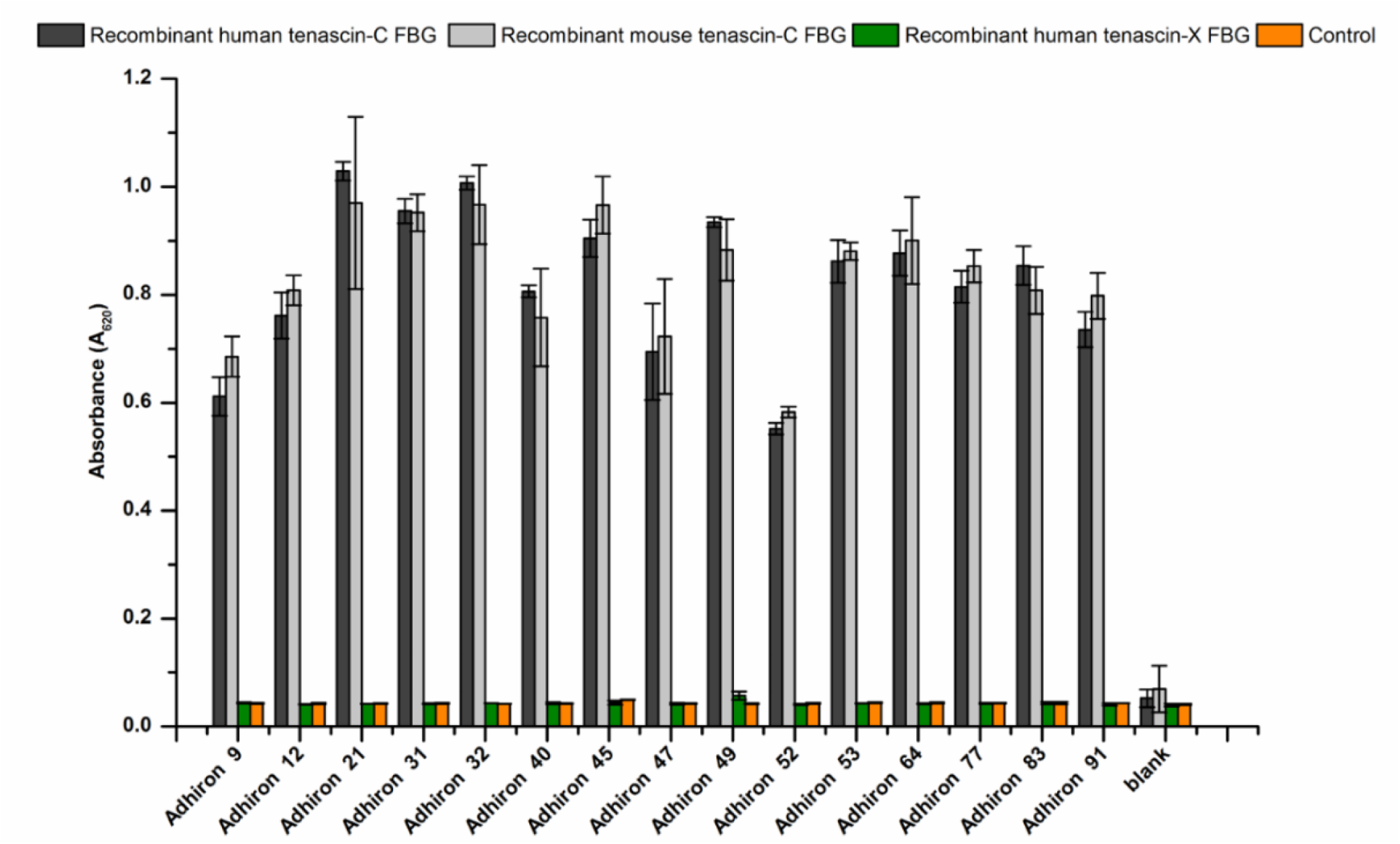
Screening selected Adhirons for cross reactivity using phage ELISA. 15 clones were tested in wells coated with recombinant human tenascin-C FBG, human tenascin-X FBG, mouse tenascin-C FBG and wells containing only buffer (control). Absorbance of oxidised TMB at 620 nm was recorded.

**Supplementary Figure 3:**
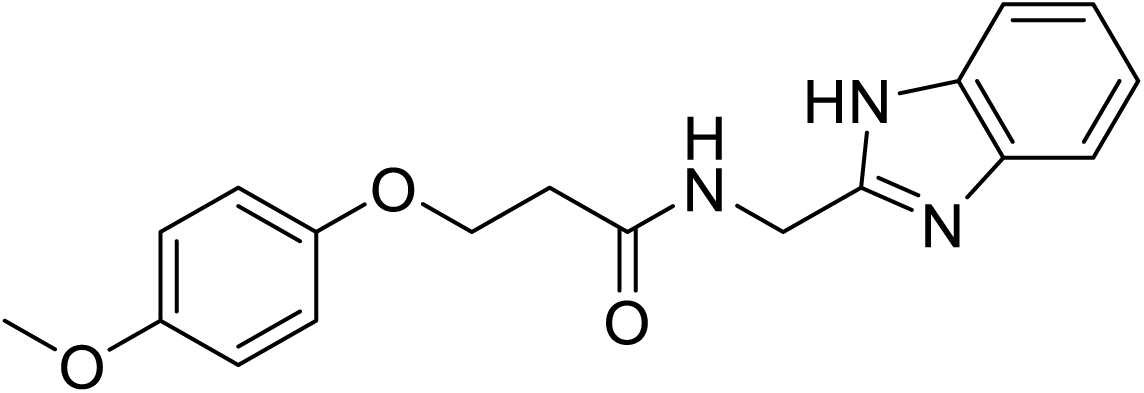
Resynthesis of Z26782404. 2-(Aminomethyl)benzimidazole dihydrochloride (75mg, 0.34 mmol) was added to a solution of 3-(4methoxyphenol)propionic acid (50mg, 0.32 mmol) in DCM (5 ml). Triethylamine (0.11 ml, 0.82 mmol) was added to the solution followed by EDC hydrochloride (125 mg, 0.63 mmol). The reaction mixture was stirred for 16 h at room temperature before being diluted with water (5 ml), extracted with EtOAc (3 x 15 ml) and the organic layer washed with brine (3 x 5 ml). The organic layer was then dried with MgSO_4_ and filtered before being concentrated under reduced pressure to give a crude product which was purified by flash column chromatography eluting with MeOH-DCM (1:49). This gave <u>the title compound</u> (24 mg, 0.11 mmol, 23%) as a pale pink solid. **M.pt.** 232.1-232.2°C; ***R_f_*** 0.7 (1:49 MeOH-DCM); **HPLC** RT = 2.30 min >99%; **δ_H_ (500 MHz, DMSO-*d_6_*);** 12.13 (1H, s), 8.57 (1H, t, *J* 5.6), 7.49-7.44 (1H, m) 7.39-7.34 (1H, m), 7.06 (2H, td, *J* 7.3 1.6), 6.81-6.71 (4H, m), 4.43 (2H, d, *J* 5.6), 4.07 (2H, t, *J* 6.3), 3.60 (3H, s), 2.56 (2H, t, *J* 6.3); **δ_C_ (125 MHz, DMSO-*d_6_*);** 170.5, 153.9, 152.9, 152.6, 143.6, 134.7, 122.3, 121.6, 118.8, 115.9, 115.0, 111.7, 64.9, 55.8, 37.6, 35.8; **ν_max_/cm^-1^ (solid)**; 3189, 3003, 2925, 2850, 1867, 1647, 1565, 1507, 1459, 1426, 1380.

---

**Supplementary Table 1:** Data collection and refinement statistics (molecular replacement)

|  | TNC/Adhiron complex |
| --- | --- |
|  | PDB 9R5Y |
| **Data collection** |  |
| Space group | P21 21 21 |
| Cell dimensions |  |
| *a*, *b*, *c* (Å) | 77.535, 90.152, 104.409 |
| $\alpha$, $\beta$, $\gamma$ (°) | 90.00, 90.00, 90.00 |
| Resolution (Å) | 58.85-1.4 (1.436-1.399) * |
| $R_{\text{sym}}$ | 0.075 (0.823) |
| $\langle I/\sigma(I) \rangle$ | 16.1 (1.8) |
| Completeness (%) | 99.6 (95.5) |
| Redundancy | 12.8 (7.5) |
| **Refinement** |  |
| Resolution (Å) | 58.85-1.40 |
| No. reflections | 143762 (704) |
| $R_{\text{work}} / R_{\text{free}}$ | 0.16/0.18 |
| No. atoms | 4968 |
| Protein | 4633 |
| Water | 338 |
| *B*-factors |  |
| Protein | 21.04 |
| Water |  |
| R.m.s. deviations |  |
| Bond lengths (Å) | 0.005 |
| Bond angles (°) | 1.295 |

\*Number of xtals for each structure should be noted in footnote. \*Values in parentheses are for highest-resolution shell.
[AU: Equations defining various *R*-values are standard and hence are no longer defined in the footnotes.]
[AU: Ramachandran statistics should be in Methods section at the end of Refinement subsection.]
[AU: Wavelength of data collection, temperature and beamline should all be in Methods section.]

**Supplementary Table 2:**
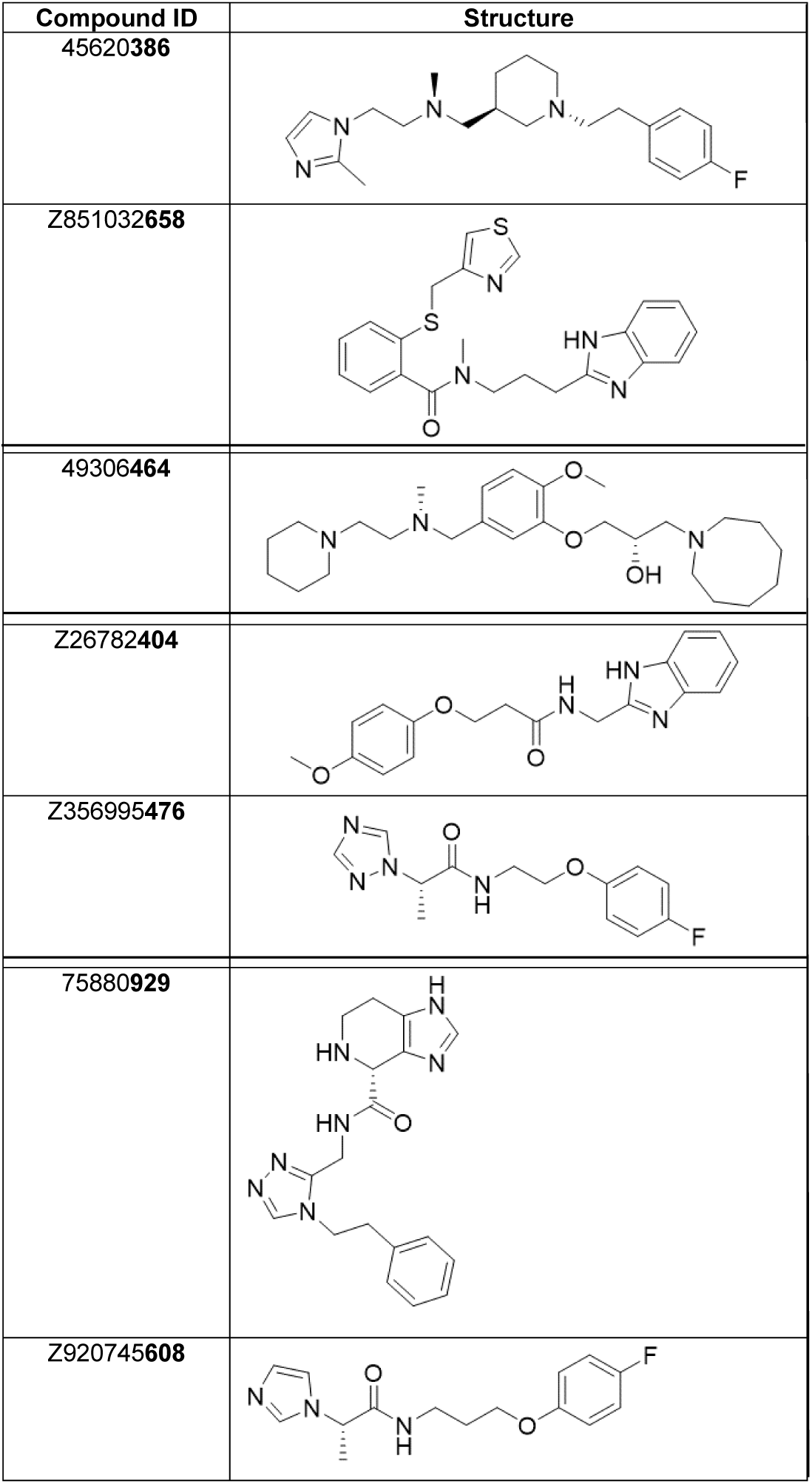
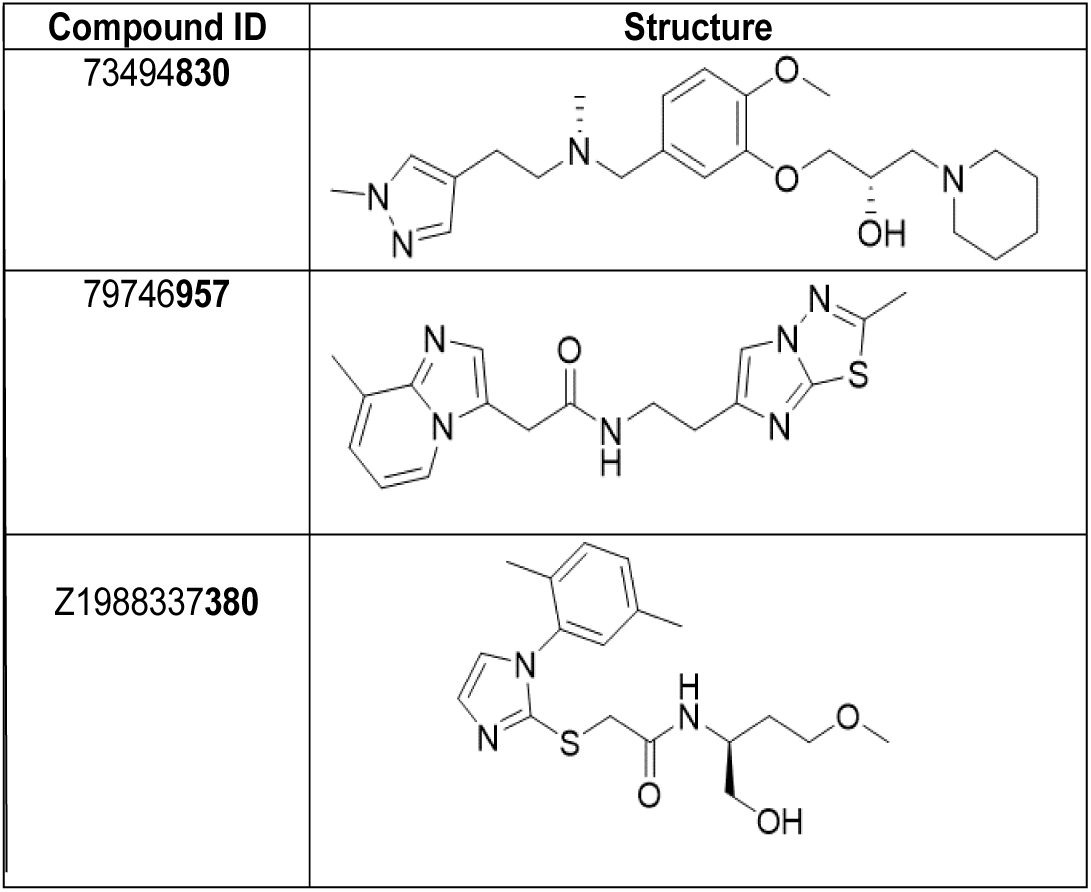
Compounds identified and purchased as mimics of Adhiron loop 1.

| Compound ID | Structure |
| --- | --- |
| 45620386 |  |
| Z851032658 |  |
| 49306464 |  |
| Z26782404 |  |
| Z356995476 |  |
| 75880929 |  |
| Z920745608 |  |
| 73494830 |  |
| 79746957 |  |
| Z1988337380 |  |

